# Familial hypercholesterolemia: a single-nucleotide variant (SNV) in mosaic at the Low density lipoprotein receptor *(LDLR)*

**DOI:** 10.1101/266874

**Authors:** Sonia Rodríguez-Nóvoa, Concepción Alonso, Carmen Rodríguez-Jiménez, Lara Rodriguez-Laguna, Gema Gordo, Victor Martinez-Glez, Iluminada García Polo

## Abstract

**Introduction:** Familial hypercholesterolemia (FH) is most frequently caused by genetic variants in the *LDLR* gene. Most of *LDLR* pathogenic variants are missense, followed by splicing and deletion/insertions variants. Mosaicism is a genetic condition in which an individual shows more than one clone of cells with different genotypes.

**Objective:** Molecular characterization of a patient with hypercholesterolemia.

**Methods:** Genetic analysis of DNA from peripheral blood and saliva was performed by NGS, sanger sequencing and pyrosequencing technologies.

**Results:** NGS analysis detected the pathogenic variant *LDLR:c.1951G>T*:p.(Asp651Tyr) in 9%-12% of reads. The presence of the variant was confirmed by pyrosequencing analysis.

**Conclusion:** Herein, we report the first case of a mosaic single nucleotide variant affecting the *LDLR* gene in a patient with familial hypercholesterolemia. As has been described for other pathologies, mosaicism could be underestimated in FH and its detection will improve with the introduction of NGS technologies in the diagnostic routine.

## Introduction

Different types of pathogenic variants in genes can cause genetic diseases. Most of them are missense variants that affect the coding region of genes. This type of variants consists in a change of aminoacid that alters or impairs the normal function of the resultant protein, therefore causing the disease phenotype. Until recently it was considered that pathogenic variants were always present in all the cells of an individual. However, it is increasingly evident that all individuals are actually complex mosaics composed of multiple genotypes acquired post-zygotically, and that in many cases this can produce a pathogenic effect. In the later cases, the variant must be present in the appropriate cell types and achieve a threshold in the number of cells affected so that the disease can be clinically detected. As a general rule, the earlier during embryonic development the mutation occurs, the greater the number of affected cells, and the greater the possibility of developing the disease. In diagnostic terms, not detecting variants in low mosaics, rather common with classical diagnostic methods, can lead to erroneous conclusions about the results of molecular tests, and therefore in the treatment and follow-up of patients ^1^.

Familial hypercholesterolemia (FH, MIM: 143890) is a genetic disorder characterized by an elevation in total cholesterol (TC) and low-density lipoproteins-cholesterol (LDL-C) leading to premature cardiovascular disease ^2^. The worldwide prevalence of FH has been estimated in 1/200-300 ^3,4^ and the main cause is the presence of pathogenic variants in the low density lipoprotein receptor gene (*LDLR*, MIM: 606945) which is involved in the LDL uptake. Up to date, more than 1700 variants have been reported in the *LDLR* gene and around 81% were classified as pathogenic or likely pathogenic ^5^. Almost all types of variants have been described in LDLR being the missense the most frequent ones. Splicing variants, small and gross deletions and insertions have been described as well.

The increase in the number of reported variants in *LDLR* is due to the implementation of screening strategies to early detection of patients with FH and to the development of new molecular strategies. Conventional method for molecular diagnosis of FH is Sanger sequencing but due to its cost and time consuming, other methodologies have been developed such as the arrays-based methods ^6^. The arrays methods are tailored to the most frequent mutations but this is a limitation because the less frequent or new variants are not detected which is an important methodology limitation. With the introduction of new technologies such as next generation sequencing (NGS), the success of the molecular diagnosis rate has increased. Despite this fact, a great number of patients remain without genetic confirmation. Traditionally, most of these patients are further classified as probably having a polygenic hypercholesterolemia.

Herein, we report a case of mosaicism in the *LDLR* gene in a patient with familial hypercholesterolemia.

## Patients and Methods

The proband is a man 57 years old referred to our reference laboratory for genetic testing in 2016 because of a clinical suspicion of familial hypercholesterolemia. He has a personal history of hypercholesterolemia detected when he was 40, with the highest levels of cholesterol documented in 2008, CT: 282 mg/dL, LDL: 195 mg/dl. There is no history of tobacco or alcohol consumption. The patient had hypertension, currently under treatment. Different pathologies such as hypothyroidism, kidney disease, diabetes, or the consumption of drugs (progestogens, anabolic steroids) were discarded. At the age of 52, he suffered acute myocardial infarction, needing percutaneous transluminal coronary angioplasty (PTCA), and implantation of a drug-eluting stent. At the age of 56, a coronary angiography showed severe stenosis in the first marginal obtuse.

As a family history, his father died at age of 77, probably due to cardiac insufficiency, the cholesterol levels are not known; his 93 years old mother is still alive, and she is not affected from dyslipidemia; and proband’s sister shows normal levels of cholesterol. The patient has four children (33, 31, 30 and 27 years old). Both younger children had hypercholesterolemia detected at the age of 5, and both were treated with resins.

Physical examination showed a height of 1.69 m, weight 74 kg, body-mass index (BMI) of 25.91; he had no arcus cornealis or xanthomas or xanthelasmas.

The patient was clinically classified as having a probably familial hypercholesterolemia according to Dutch Lipid Clinic Network Criteria (DLCNC), so the genetic study of familial hypercholesterolemia was performed.

Genetic analysis was performed by NGS using a customized panel of 198 genes. Preparation and exome enrichment steps were performed according to manufacturer’s workflow (Roche Nimblegen) and it was sequenced using MiSeq system. A subset of genes was chosen for analysis of hypercholesterolemia: *LDLR*, *APOB*, *PCSK9*, and *LDLRAP1*. Bioinformatic analysis was performed using algorithms developed by our bioinformatics unit. The *in silico* predictors of pathogenicity used were CADD (Combined Annotation Dependent Depletion), Polyphen (Polymorphism Phenotyping), MutAssesor, Fasthmm, and Vest. Scores of conservation used were Gerp2, PhasCons, Phylop. The splicing predictors used were MaxEntScan, NNSplice, GeneSplicer and Human Splicing Finder. Variants with minor allele frequency >1% were excluded from further analysis. The files were uploaded in BAM format for analysis using Alamut software.

Multiplex ligation-dependent probe amplification (MRC-Holland) was used for detection of large insertions/deletions in *LDLR* gene.

Sanger sequencing was used to confirm variants present in more than 15% of the reads in the deep sequencing data: Standard PCR and Sanger sequencing were performed using the 96-capillary ABI 3730xl ADN analyzer (Applied Biosystem, Foster, USA). The validation of variants detected in mosaics over 5% of the readings in the NGS was performed by the pyrosequencing technique: primers were designed using PyroMark software, and QIAGEN reagents and the Pyromark Q96 MD instrument (QIAGEN, USA) were used according to manufacturer’s protocol.

## Results

NGS data from the proband’s DNA blood sample was suitable for analysis after passing the quality parameters established in our laboratory: number of reads more than 30x in the 99% of the target bases. The results of the bioinformatics analysis of data from NGS showed the *c.1951G>T*:p.(Asp651Tyr) variant in 12% of the reads, located in exon 13 of the *LDLR* gene. This is a known pathogenic variant associated with FH ^7^ Sanger sequencing was negative for this variant. The presence of large deletions or insertions in the whole gene was discarded by MLPA.

When we performed a visual analysis of the bam file using Alamut software we observed three types of reads within exon 13. Close to our pathogenic variant of interest (*c.1951G>T*) we found the synonymous polymorphism c.1959T> C (rs5925), with a MAF / MinorAlleleCount for C = 0.336 / 1682 according to dbSNP. This allowed us the visualization of three different compound genotypes: the first one included the pathogenic variant *c.1951G>T* plus the wild type allele for position c.1959T in approximately 12% of the reads. The second one carries the wild type allele for position c.1951G and the polymorphism c.1959T>C in 48% of the reads; and the third one carries the wild type allele for position c.1951G and the c.1959T in 40% of the reads (**Figure 1**).

**Figure 1.**
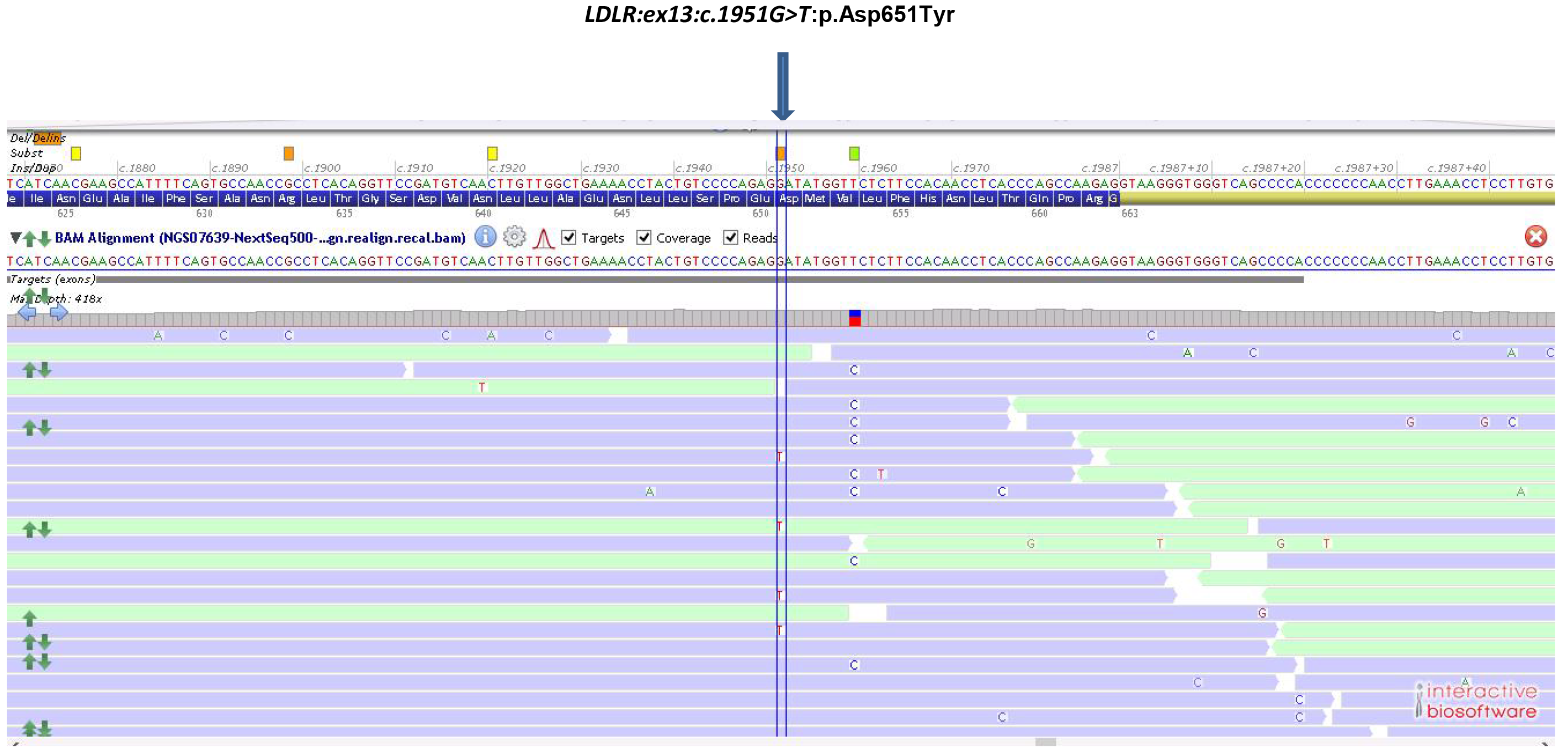
NGS analysis of proband’s blood. Visualization of BAM file using Alamut software focus on the exon 13 of *LDLR*. The arrow indicates the position of the genetic variant *LDLR:ex13:c.1951G>T*:p.(Asp651Tyr).

A new blood sample from the proband was requested in order to confirm the results. After DNA extraction and NGS analysis of the new sample, the pathogenic variant *c.1951G>T* was observed in 9% of reads, confirming the previous results.

The variant *c.1951G>T*:p.(Asp651Tyr) has been previously described and associated with familial hypercholesterolemia ^7^. It affects a highly conserved nucleotide (phyloP: 5.69 [-14.1;6.4]) and a highly conserved amino acid (considering 11 species). There is also a large physicochemical difference between Asp and Tyr (Grantham dist.: 160 [0-215]). We use the *in silico* predictors of pathogenicity CADD, VEST, Sift, and Mutation taster and all of them classified the variant as deleterious. This variant is located at the protein domain LDLR class B repeat. Align GVGD: C15 (GV: 85.08-GD: 92.25). According to AMCG criteria for genetic variant classification, this variant was classified as pathogenic.

We extended the family study by NGS to the proband’s two children with hypercholesterolemia. Both of them were heterozygous for the variant *c.1951G>T*:p.(Asp651Tyr). These results were confirmed by Sanger sequencing. The proband’s wife and the other two children with normal levels of cholesterol showed the common allele at the position *c.1951* by Sanger sequencing. Family pedigree is shown in **Figure 2**.

**Figure 2.**
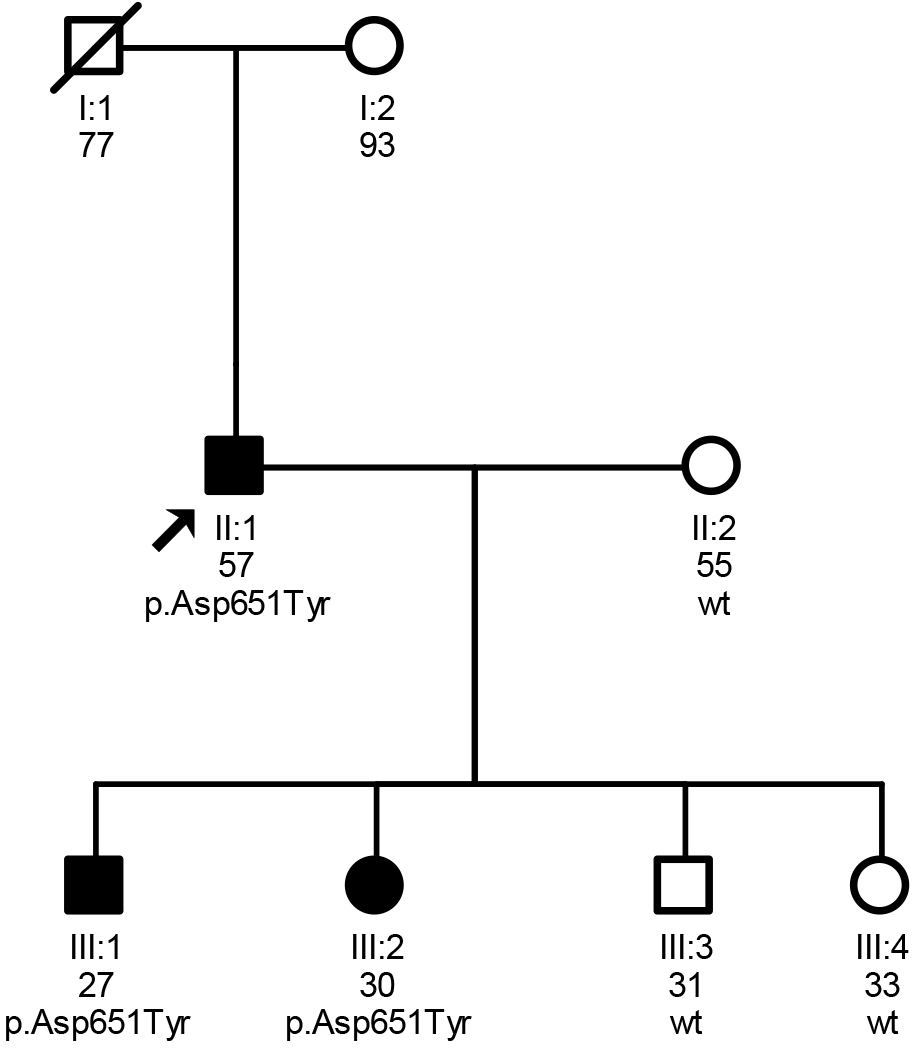
Pedigree of the family showing the segregation of the variant *LDLR:ex13:c.1951G>T*:p.(Asp651Tyr). The arrow indicates the proband individual; circle and square symbols represent women and men respectively; shadow filled symbols indicate the affected members with hypercholesterolemia. Line 1 below symbols correspond to the individual identification, line two indicates the age; line 3 indicates the LDLR genotype: p.(Asp651Tyr) or normal genotype at this position (wt).

In order to confirm the mosaicism at position c. *1951* of the *LDLR* by an orthogonal method, we performed **pyrosequencing** analysis in the DNA extracted from peripheral blood from the proband, using as a positive control the older daughter, who was heterozygous for the variant *c.1951G>T* by NGS. The results showed that the proband is a mosaic with the pathogenic variant found in 20% of the reads in peripheral blood DNA. Heterozygous state was confirmed in the daughter. Since mosaicism can affect different tissues in a different proportion, we also decide to analyze a **saliva** sample from the proband, using the same three technologies. NGS sequencing and pyrosequencing analysis showed a mosaic of 8% and 18% of the pathogenic variant respectively, whereas Sanger sequencing hardly showed the variant. **Figure 3** summarizes the results of blood and saliva at the position *c.1951* at the exon 13 of the *LDLR* using the different types of technologies.

**Figure 3.**
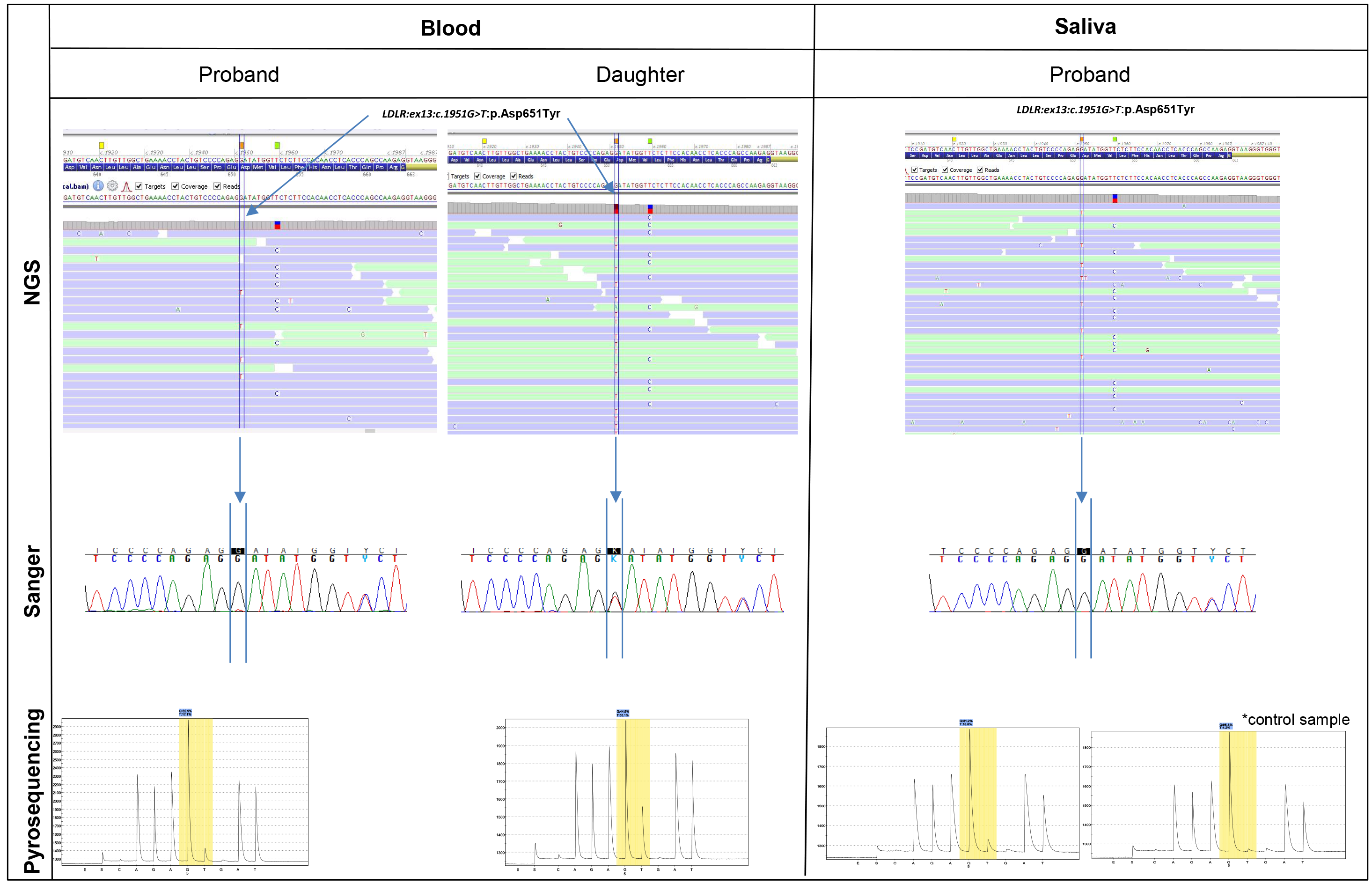
The figure shows the results of NGS, Sanger sequencing and pyrosequencing analysis performed in blood and saliva samples. NGS analysis of blood samples show the reads with the pathogenic variant in the proband and his daughter. Alamut software was used to visualize BAM file. Sanger sequencing corresponding to exon 13 of the *LDLR* shows that the variant is hardly detected in the proband whereas it is clearly detected in his daughter. Pyrosequencing analysis of proband and daughter blood shows approximately 20% and 50% of pathogenic variant, respectively. Saliva sample from the proband was analyzed using the same three technologies. NGS sequencing and pyrosequencing analysis showed a mosaic of 8% and 18% of the pathogenic variant respectively, whereas Sanger sequencing hardly shows the variant.

## Discussion

In this study we report the first case of a mosaic single-nucleotide variant (SNV) affecting the most relevant gene involved in familial hypercholesterolemia, the *LDLR* gene. This phenomenon occurs when a *de novo* genetic variant appears post-zygotically causing a mosaicism in the embryo. The timing and location in which the variant occurs plays a key role in the distribution of the variant among tissues (that is, somatic, germline or gonosomal) and therefore, determine the final phenotype in the affected patient as well as the patterns of disease reoccurrence within families. Several biological specimens can be tested to detect mosaicism. In this study, we have analyzed two types of specimens, blood and saliva. Although the liver is the target tissue for molecular diagnosis of hypercholesterolemia, since it is the main organ responsible for lipid homeostasis, it is not justified to obtain such a type of sample for genetic analyses due to the invasive nature of sampling. In any case, we assert that the proband’s liver is affected to such an extent grade conferring the patient with hypercholesterolemia phenotype. The proband’s germline tissue is also affected since two of his children are carriers of the pathogenic variant in a heterozygous form.

The presence of mutations in the form of somatic mosaicism is well known in pathologies that, in a classical way, are caused by constitutionally dominant mutations, and therefore compatible with life. As an example, the neurofibromatosis type I due to mutations in one of the most extensive genes in humans, the *NF1* gene. However, mutations in this gene can also occur in the form of somatic mosaicism giving rise to what is known as segmental NF1, since the clinical manifestations are limited to a part of the body ^8^. Other examples of dominant diseases that may occur as mosaic include Duchenne myotonic dystrophy and the hereditary haemorrhagic telangiectasia.

Other pathologies present only in the form of mosaicism and are not heritable, probably because they are incompatible with life in germinal form. As an example we have the *PIK3CA* related overgrowth spectrum (PROS) ^9^ or the McCune-Albright syndrome. A third form of mosaicism is the one causing germline mosaic disorders, in which dominant pathologies occur in several children of phenotypically healthy parents, due to the presence of a mutation in the germline tissue of one of the parents. The most reported pathology with this type of mechanism is osteogenesis imperfecta type II ^8^.

The mosaic variant in the *LDLR* gene presented in this work belongs to the group of mosaicisms in dominant genes. However, in addition to the somatic mosaicism, the patient transmitted the variant to two of his children, so his germ cells must be affected. This combination between somatic mosaicism and germinal mosaicism is known as gonosomal mosaicism.

The case described here confirms and highlights the fact that mosaicism does not always occur in clinically easily identifiable disorders. Therefore, we consider that in the FH the molecular confirmation of a mosaic pathogenic variant is especially useful in those patients with LDL-c borderline, and allows cascade screening in the family to further establish lifestyle changes or medication for early cardiovascular prevention. Although heterozygous FH frequency is around 1/200-250, a minor fraction of them is identified ^10^, for this reason improving the molecular detection is of great interest. The use of new technologies such as the high-throughput deep sequencing allows the detection of mosaics. As we describe in our study it is possible that mosaicism can explain some of the cases of patients without molecular confirmation. This phenomenon can be suspected specially in those molecular unconfirmed cases with strong familial segregation, even more when less sensitive methods for analysis were previously used. Even if NGS was used for molecular diagnosis of FH, the bioinformatic treatment of the data should consider the possibility of a mosaicism so that these types of variants are not filtered prior to the variant calling. For example, in routine laboratories, a common variant call ratio used for filtering variants leaves out those that not reach the 20% of the reads. Therefore, the bioinformatics algorithms used for routine analysis need to improve in order to better detect mosaicism. Recent publications have addressed this topic ^11^. On the other hand, another aspect to take into account is the reading depth usually used. Improving the deep coverage up to 1000× will increase the ability to detect low-level mosaicism. However, at present a relatively high cost of the test may be a major limitation for large-scale application of this approach in diagnostic laboratory. Something that will probably be solved soon with the technological advances in the NGS platforms.

Sanger sequencing has been the gold standard technology used in most laboratories but it fails to identify cases of mosaicism. It is difficult to detect when the grade of mosaicism is low. Sanger sequencing possess less sensitivity compared with massive sequencing in detecting mosaicism. Recent publications state that the use of sanger sequencing is not adequate to validate NGS findings specially when the last one shows good quality parameters such as a good coverage of reads and quality ^12–14^. The authors state that the use of sanger validation should be restricted for those cases in which the result of NGS has a serious implication for the probands and their families ^12^.

With the extensive use of massive sequencing techniques more genes are screened routinely providing ever more information. With adequate tools to process the information, it is possible to explore not only the main genes responsible for FH but other candidate genes. In our experience, certain patients remitted to our laboratory because suspicious of FH have genetic variants in other genes involved in cholesterol metabolism. It is possible that these candidate genes could explain the lipid disturbances in these patients, and mosaicism could also be present in them.

Nowadays, there is no doubt that next generation sequencing provides us with the better results, improving not only the detection of new mutations, but also allowing the identification of new genes and mosaicism. Further analysis is warranted to know the extent of the phenomenon of mosaicism not only in familial hypercholesterolemia but in other dyslipemias.

## Author contributions

SRN, analyzed the NGS data, design the study and write the manuscript; CA and IGP contributed to clinical management, and follow-up of the patient and review the manuscript; CRJ performed sanger analysis and figures; LRL and GG performed pyrosequencing analysis and figures; VM-G contributed to review the data and the manuscript. All authors have approved the final article.

